# Microscopic and metatranscriptomic analyses revealed unique cross-domain symbiosis between *Candidatus* Patescibacteria/candidate phyla radiation (CPR) and methanogenic archaea in anaerobic ecosystems

**DOI:** 10.1101/2023.08.25.554742

**Authors:** Kyohei Kuroda, Meri Nakajima, Ryosuke Nakai, Yuga Hirakata, Shuka Kagemasa, Kengo Kubota, Taro Q.P. Noguchi, Kyosuke Yamamoto, Hisashi Satoh, Masaru K. Nobu, Takashi Narihiro

## Abstract

To verify the parasitic lifestyle of *Candidatus* Patescibacteria in the enrichment cultures derived from a methanogenic bioreactor, we applied multifaceted approaches combining cultivation, microscopy, metatranscriptomic, and protein structure prediction analyses. Cultivation experiments with the addition of exogenous methanogenic archaea with acetate, amino acids, and nucleoside monophosphates and 16S rRNA gene sequencing confirmed the increase in the relative abundance of *Ca*. Patescibacteria and methanogens. The predominant *Ca*. Patescibacteria were *Ca*. Yanofskybacteria and 32-520 lineages (to which belongs to class *Ca*. Paceibacteria) and positive linear relationships (*r*^*2*^ ≥ 0.70) between the relative abundance of *Ca*. Yanofskybacteria and *Methanothrix*, suggesting that the tendency of the growth rate is similar to that of the host. By fluorescence *in situ* hybridization (FISH) observations, the FISH signals of *Methanothrix* and *Methanospirillum* cells with *Ca*. Yanofskybacteria and with 32-520 lineages, respectively, were significantly lower than those of the methanogens without *Ca*. Patescibacteria, suggesting their parasitic interaction. The TEM and SEM observations also support parasitism in that the cell walls and plugs of these methanogens associated with submicron cells were often deformed. In particular, some *Methanothrix*-like filamentous cells were dented where the submicron cells were attached. Metatranscriptomic and protein structure prediction analyses identified highly expressed secreted genes from the genomes of *Ca*. Yanofskybacteria and 32-520, and these genes contain adhesion-related domains to the host cells. Considering the results through the combination of microscopic observations, gene expression, and computational protein modeling, we propose that the interactions between *Ca*. Yanofskybacteria and 32-520 belonging to class *Ca*. Paceibacteria and methanogenic archaea are parasitism.

## Main text

Candidate phyla radiation (CPR)/*Candidatus* Patescibacteria, ultrasmall bacteria, is a major lineage of the domain Bacteria that is widely distributed in various natural and artificial environments (1–5). To date, several *Ca*. Patescibacteria-bacteria intradomain symbioses have been observed (*e*.*g*., *Ca*. Saccharimonadia with Actinobacteria (6, 7) and *Ca*. Gracilibacteria with Gammaproteobacteria (8, 9). Most recently, three cases of cross-domain symbioses between the class *Ca*. Paceibacteria (former Parcubacteria/OD1) of the *Ca*. Patescibacteria and domain Archaea have been reported by using cultivation and microscopic observations, *i*.*e*., *Ca*. Yanofskybacteria/UBA5738 (10) and *Ca*. Nealsonbacteria (11) with aceticlastic methanogenic archaeon *Methanothrix* and 32-520/UBA5633 with the hydrogenotrophic methanogen *Methanospirillum* (12). Our previous microscopic observations suggested that the interactions between *Ca*. Paceibacteria and methanogenic archaea are likely to cause parasitism, based on the absence or low detectable ribosomal activity (based on fluorescence *in situ* hybridization [FISH]) and deformations at the attachment sites (based on transmission electron microscopy [TEM]) (10, 12). In addition, several genetic features that may contribute to parasitism have been identified in the metagenome-assembled genomes of *Ca*. Paceibacteria (10–12); however, the challenge is to maintain enrichment cultures to ensure that those genes are expressed. In the present study, to uncover the underlying mechanisms of parasitic interactions between *Ca*. Paceibacteria and methanogens, we attempted to perform a multifaceted approach combining cultivation and microscopy along with metatranscriptomic analyses for enrichment cultures derived from anaerobic bioreactors.

To set up the experimental design for these analyses, we prepared seven parallel enrichment cultures (called C-1–C-7) transferred from the culture system C-d2-d1 (10), which contains acetate, amino acids, and nucleoside monophosphates as potential growth factors for *Ca*. Patescibacteria (see Text S1). The enrichment cultures showed the production of methane gas on Days 14 and 31. We then analyzed the microbial community structures by using 16S rRNA gene amplicon sequencing on Days 7, 14, 21, and 31. The abundances of *Ca*. Yanofskybacteria OTU0011 (PMX_810_sub as the metagenomic bin), 32-520 OTU 0014 (PMX.108), and 32-520 OTU0072 (PMX.50) (Fig. S1A and S1B) during cultivation varied from 0.15–12.5%, 0.6–2.3%, and 0.1–0.87%, respectively (Fig. S2A–S2D and Text S1).

The physiological and morphological characteristics of the symbioses were confirmed by microscopic observations based on fluorescence *in situ* hybridization (FISH), TEM, and scanning electron microscopy (SEM). On Day 31, the FISH fluorescence of *Methanothrix* filaments with more than 5 *Ca*. Yanofskybacteria cells was significantly lower than that of *Methanothrix* cells without *Ca*. Yanofskybacteria cells because of the significantly larger areas with no fluorescence (Fig. 1A, *p* < 0.05). In addition, the fluorescence fractions (clear, weak, and no fluorescence) of the *Methanothrix* filaments also showed that many of the *Ca*. Yanofskybacterial cells (35±25 cells/*Methanothrix*-filamentous) were attached to *Methanothrix* with no fluorescence on Day 31 (Fig. 1B and 1D, Figs. S3A, and S4). *Methanospirillum* cells with 32-520 cells (1.1±0.3–1.3±0.5 cells/*Methanospirillum*-cell, Fig. S3B) also had significantly lower FISH signals than *Methanospirillum* cells without 32-520 (Fig. 1C and Fig. S5, *p* < 0.05). Taken together, the interactions between methanogenic archaea and *Ca*. Paceibacteria are parasitic, strongly supporting previous predictions with statistical evidence (10, 12). The TEM observations also supported the parasitism of *Ca*. Yanofskybacteria, as the cell walls of *Methanothrix* (sheathed filamentous cells) (13) were often deformed where the submicron cells were attached. Interestingly, some *Methanothrix*-like filamentous cells with submicron *Ca*. Yanofskybacteria-like cells were dented through TEM observation (Fig. 1E–1H). In addition, the submicron cells produced adhesive materials at the attachment sites on the *Methanothrix* cells (Fig. 2A and 2B). Therefore, it is speculated that the secreted materials are important for the attachment of *Ca*. Yanofskybacteria cells to *Methanothrix* cells. Another type of submicron cell (likely 32-520 cells) was tightly attached to the plug structures of *Methanospirillum* (rod-shaped sheath cells) (Fig. 2C and 2D) (14). The TEM observations indicated that there are extracellular substances at the attachment sites of 32-520-like submicron cells (Fig. 1I and 1J). These microscopic observations imply that the production of extracellular substances is essential for the episymbiosis of *Ca*. Paceibacteria. High-resolution imaging techniques such as cryo-electron microscopy can be an effective approach to further clarify their cellular structures and attachment sites.

**Figure 1.**
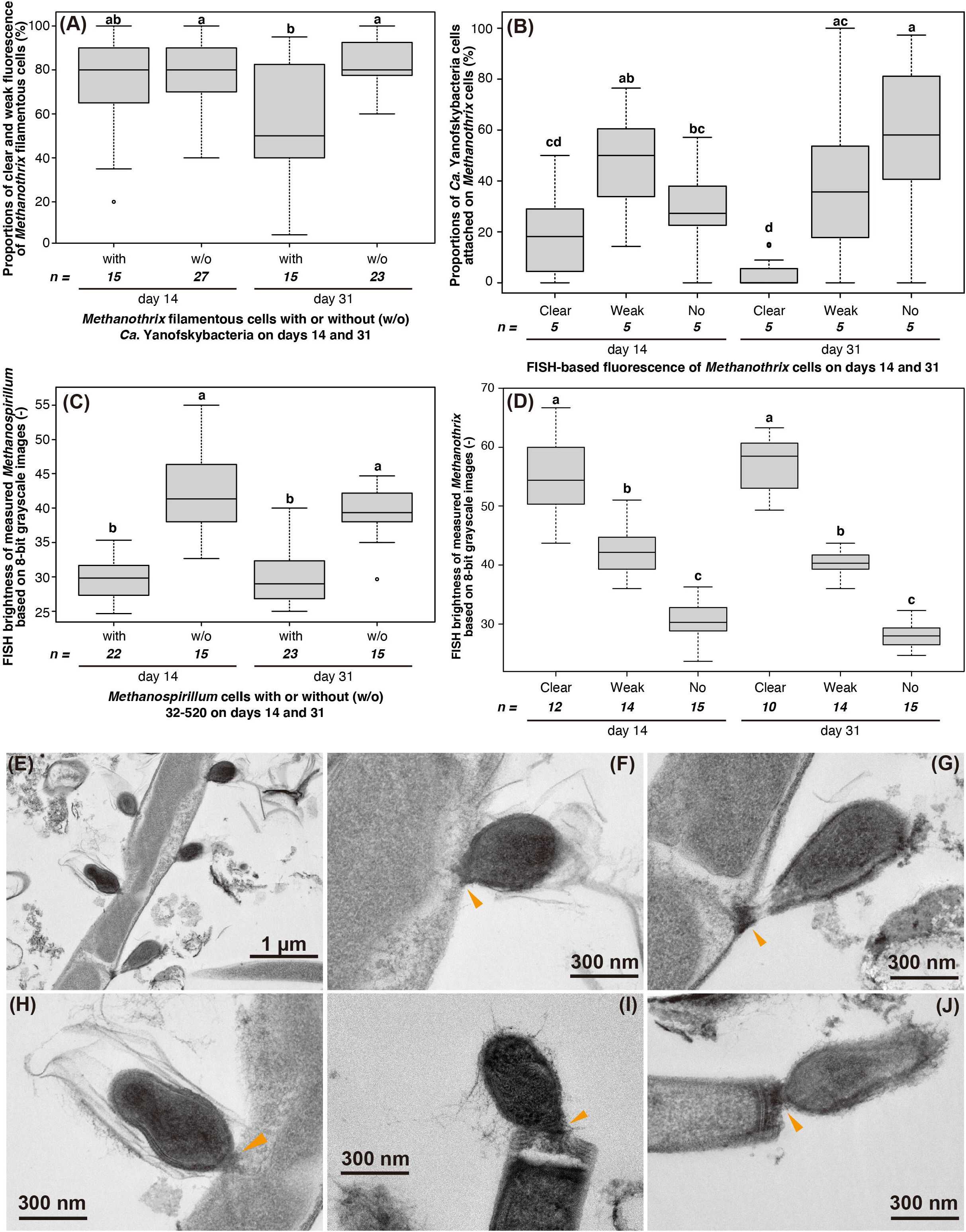
(A) Cell length proportions of clear and weak fluorescence of *Methanothrix* filamentous cells calculated based on the fluorescence *in situ* hybridization signals using the *Methanothrix*-targeting MX825-FITC probe and *Candidatus* Yanofskybacteria-targeting Pac_683-Cy3 probe. The *Methanothrix* cells attached with > 5 *Ca*. Yanofskybacteria cells were chosen for calculation. (B) Proportions of detected *Ca*. Yanofskybacteria cells attached to the different fluorescence of *Methanothrix* cells: attached to *Methanothrix* filamentous cells with clear fluorescence, attached to *Methanothrix* with weak fluorescence, and attached to *Methanothrix* with no fluorescence. (C) The fluorescence of *Methanospirillum* cells with or without 32-520 cells. (D) FISH brightness of measured *Methanothrix* filamentous cells based on 8-bit grayscale images. (A)–(D) Different letters in the figure indicate significant differences among the values of the proportions based on Tukey’s test (*p* < 0.05). (E)–(I) Transmission electron micrographs of small submicron cells attached to (E)–(H) *Methanothrix*-like cells and (I) and (J) *Methanospirillum*-like cells in culture system C-1 on Day 33. Orange arrows indicate extracellular substances at the attachment sites.

**Figure 2.**
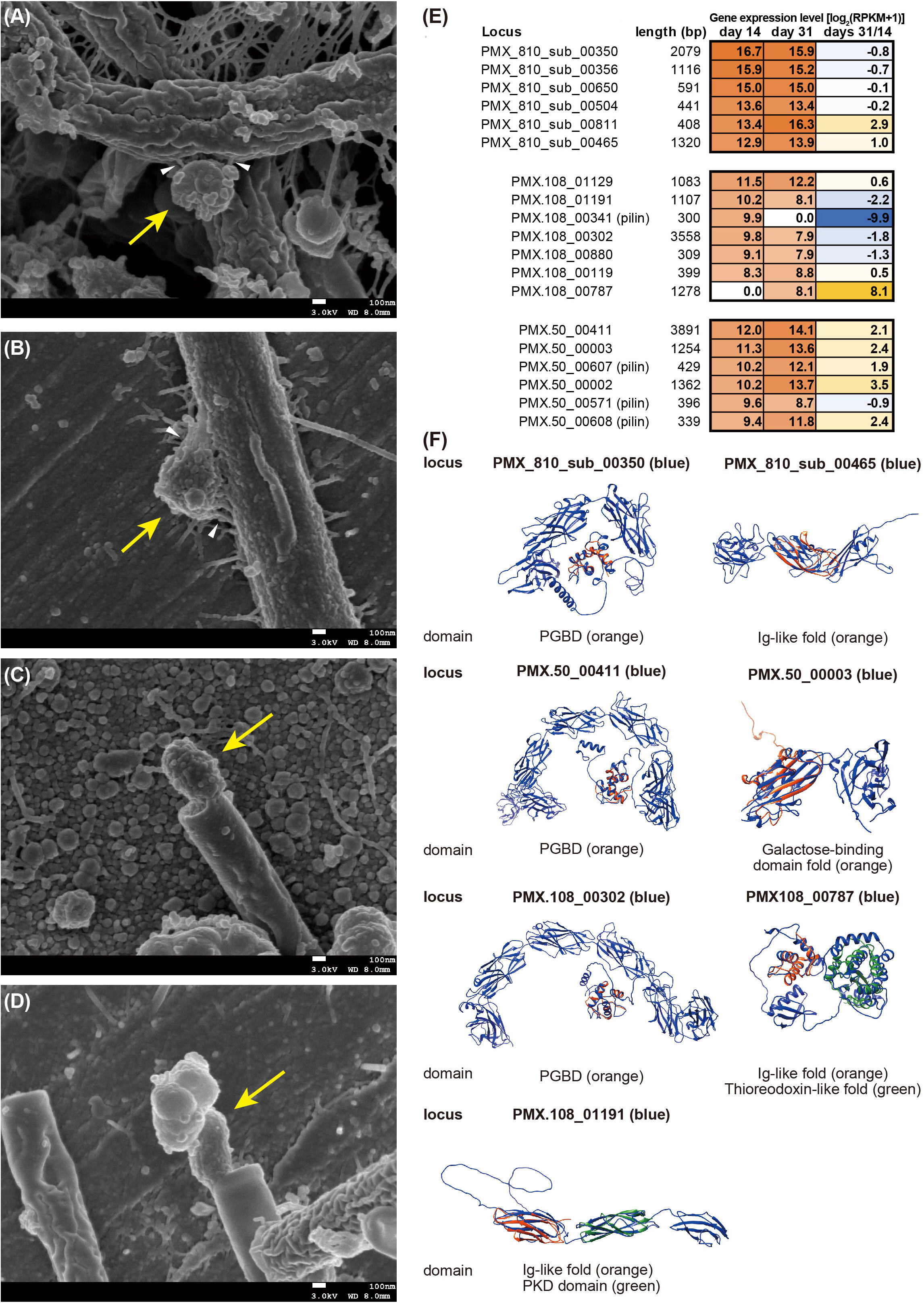
(A)–(D) Scanning electron micrographs of small submicron cells (yellow arrows) attached to (A) and (B) *Methanothrix*-like cells and (C) and (D) *Methanospirillum*-like cells in culture system C-1 on Day 33. White arrows indicate extracellular substances at the attachment sites. (E) Gene expression heatmap of the five most highly expressed genes with signal peptides in the genome of *Candidatus* Yanofskybacteria/UBA5738 (PMX.810_sub) and 32-520/UBA5633 (PMX.50 and PMX.108) in culture systems C-2–C-4 on Day 14 and C-6 and C-7 on Day 31. The color scale from white to orange shows the gene expression level based on the normalized RPKM value (see Text S1). “Days 31/14” indicates the difference in gene expression between Days 31 and 14. (F) Predicted protein structures of the highly expressed genes of *Ca*. Patescibacteria using the AlphaFold2 software package (24). The overlaying domain was predicted through the InterPro database (http://www.ebi.ac.uk/interpro/). PGBD, Ig-like, and PKD are the peptidoglycan binding domain, immunoglobulin-like domain, and polycystic kidney disease domain, respectively.

To confirm their interactions based on gene expression levels, we performed metatranscriptomics for the enrichment cultures on Days 14 (triplicate) and 31 (duplicate). A total of 6.0–10.8 Gb sequences were obtained and mapped to the previously reconstructed metagenome-assembled bins of *Ca*. Yanofskybacteria/UBA5738 (PMX_810_sub) and 32-520/UBA5633 (PMX.108 and PMX.50), which belonged to MWCK01 and UBA5633 at the genus level in the GTDB database (15), respectively (Fig. S1B and Text S1) (10)(12). Previous studies have suggested that the competence protein ComEC, secretion systems, pilus, and several transporters are important for the symbiosis of ultrasmall microbes, including *Ca*. Patescibacteria, with hosts (16–19). Accordingly, these genes were highly expressed in the genome of *Ca*. Yanofskybacteria PMX_810_sub on Day 14 (Table S4), suggesting their importance in symbiotic lifestyles during the early growth stage. In addition, F-type H^+^-transporting ATPase proteins were highly expressed in *Ca*. Yanofskybacteria and 32-520 PMX.50 (Tables S2 and S4), which are encoded by type IV pilus assembly proteins (Table S2). In a previous study, ATPase and type IV pili were predicted to function in attachment and motility on larger host surfaces (19). Furthermore, some active peptidase-like proteins with signal peptides (PMX_810_sub_00385, PMX_810_sub_00508, PMX.108_00125, PMX.108_00310, PMX.108_00457, PMX.108_00476, and PMX.50_00413) (Table S2) and substrate-binding proteins of amino acid/metal transport systems were found in the three *Ca*. Patescibacteria genomes (Table S4). Although the detailed functions remain unclear, the addition of external sources of amino acids and trace elements may be key factors for the successful enrichment of *Ca*. Patescibacteria.

To estimate the function of the proteins encoded by the five most highly expressed but functionally unknown genes with signal peptides, including peptidoglycan binding domains (PGBD: PMX_810_sub_00350, PMX.50_00411, and PMX.108_00302), immunoglobulin (Ig)-like folds (PMX_810_sub_00465, PMX108_00787, and PMX.108_01191), galactose-binding domain folds (PMX.50_00003), thioredoxin-like domains (PMX.108_00787), polycystic kidney disease (PKD) domains (PMX.108_01191), and type IV secretion system pilins (PMX.108_00341, PMX.50_00571, PMX.50_00607, and PMX.50_00608), the protein structures were predicted computationally (Fig. 2F and Table S2). These domains are known to be host adhesion-related proteins, such as membrane-anchored secreted proteins that bind membrane substrates (20–22). Of these, PGBD is found at the N- or C-terminus of several enzymes involved in cell wall degradation (*e*.*g*., membrane-bound lytic murein transglycosylase B and zinc-containing D-alanyl-D-alanine-cleaving carboxypeptidase) (23). Interestingly, the PGBD-containing enzymes showed relatively similar structures among the three *Ca*. Patescibacteria (Fig. 2F), suggesting that these secreted uncharacterized proteins are likely to be specific and important for *Ca*. Patescibacteria-methanogen interactions.

In summary, we found that the interactions between the class *Ca*. Paceibacteria of *Ca*. Patescibacteria and methanogenic archaea are parasitism through the combination of FISH, TEM, and SEM observations and the first successful gene expression analysis of class *Ca*. Paceibacteria. In addition, we identified highly expressed secreted proteins with PGBD that have similar structures among three *Ca*. Paceibacteria. The microscopic and metatranscriptomic observations suggested that the adhesion/degradation functions of *Ca*. Paceibacteria to host methanogen cells are uniquely developed for their parasitic lifestyle. Further elucidation of the characterizations of the cell[cell interactions in detail and the establishment of refined cocultures of *Ca*. Paceibacteria and methanogens are essential to clarify the influence of ultrasmall bacteria on anaerobic ecosystems.

## Acknowledgments

This study was partly supported by the Japan Society for the Promotion of Science KAKENHI JP16H07403 and JP21H01471, a matching fund between the National Institute of Advanced Industrial Science and Technology (AIST) and Tohoku University, and research grants from the Institute for Fermentation, Osaka (G-2019-1-052 and G-2022-1-014). The authors thank Riho Tokizawa, Yuki Ebara, Tomoya Ikarashi, and Maho Takai at AIST for technical assistance.

## Contributions

K. Kuroda and T.N. designed this study. K. Kuroda and M.N. performed sampling, cultivation, microscopy, and sequence analysis. K. Kuroda, M.N., R.N., Y.H., S.K., K. Kubota, T.Q.P.N., K.Y., H.S., M.K.N., and T.N. interpreted the data. K. Kuroda, M.N., and T.N. wrote the manuscript with input from all coauthors. All authors have read and approved the manuscript submission.

## Legends of Supplemental materials

### Supporting Information

The file containing materials and methods and results and discussion.

**Figure S1** Phylogenetic trees of order *Candidatus* Paceibacterales based on (A) 16S rRNA gene sequences and (B) concatenated phylogenetic marker genes of GTDBtk 2.0.0 (ver. r207) (15). The phylogenetic positions of the metagenomic bins PMX_810 and PMX.108/PMX.50 are shown in pink and blue, respectively. The 16S rRNA gene-based tree was constructed using the neighbor-joining method. Sequences that match the Pac_683 and 32-520-1066 probes are shown in yellow and blue layers, respectively.

**Figure S2** (A) Relative abundance of predominant *Candidatus* Patescibacteria and methanogenic archaea in the culture systems based on 16S rRNA gene sequencing. (B)–(D) Linear regression analysis between predominant methanogens and *Ca*. Patescibacteria based on 16S rRNA gene-based relative abundance. (B) *Methanothrix* OTU0004 and *Ca*. Yanofskybacteria OTU0011, (C) *Methanospirillum* OTU0025 and 32-520 OTU0014, and (D) *Methanospirillum* OTU0025 and 32-520 OTU0072.

**Figure S3** (A) Number of *Candidatus* Yanofskybacteria cells attached to one *Methanothrix* filamentous cell on Days 14 and 31. (B) Number of 32-520 cells attached to one *Methanospirillum* cell on Days 14 and 31. The statistical analysis was performed based on Welch’s t test.

**Figure S4** Micrographs of (A) and (E) phase-contrast, (B) and (F) 4’,6-diamidino-2-phenylindole dihydrochloride staining, (C), (D), (G), and (H) fluorescence *in situ* hybridization obtained from the culture system C-1 on (A)–(D) Days 14 and (G)–(H) 31. (C) and (G) *Ca*. Yanofskybacteria-targeting Pac_683-Cy3 probe and (D) and (H) *Methanothrix*-targeting MX825-FITC probe.

**Figure S5** Micrographs of (A) and (E) phase-contrast, (B) and (F) 4’,6-diamidino-2-phenylindole dihydrochloride (DAPI) staining, (C), (D), (G), and (H) fluorescence *in situ* hybridization obtained from the culture system C-1 on (A)–(D) Days 14 and (G)–(H) 31. (C) and (G) 32-520-targeting 32-520-1066-Cy3 probe and (D) and (H) Archaea-targeting ARC915-FITC probe. Yellow arrows indicate FISH-detectable 32-520 cells. Light blue arrows indicate unspecific FISH signals that were not observed by phase-contrast and DAPI staining. Dashed white lines indicate weak or no FISH signals of *Methanospirillum*-like cells.

**Table S1** Mapped and total sequence reads of the metatranscriptome in this study.

**Table S2** Summary of the gene expression level and annotation of metagenomic bins of *Candidatus* Yanofskybacteria/UBA5738 (PMX_810_sub) and 32-520/UBA5633 (PMX.50 and PMX.108) using DRAM, BlastKOALA, and SignalP annotation software.

**Table S3** Summary of the gene expression level and annotation of metagenomic bins of *Methanothrix* (PMX.81, PMX.12, and PMX.35) and *Methanospirillum* (PMX.141_sub) using DRAM, BlastKOALA, and SignalP annotation software.

**Table S4** Summary of the annotation of the secretion systems, transporter-related proteins, ATPase, and cell growth-related genes in the genome of *Candidatus* Yanofskybacteria/UBA5738 (PMX.810_sub) and 32-520/UBA5633 (PMX.50 and PMX.108) in culture systems C-2–C-4 on Day 14 and C-6 and C-7 on Day 31.

