## Supporting Information for "Microscopic and metatranscriptomic analyses revealed unique cross-domain symbiosis between *Candidatus* Patescibacteria/candidate phyla radiation (CPR) and methanogenic archaea in anaerobic ecosystems"

Kyohei Kuroda[1*¶], Meri Nakajima[1,2¶], Ryosuke Nakai[1], Yuga Hirakata[3], Shuka Kagemasa[4,5], Kengo Kubota[5,6], Taro Q.P. Noguchi[7], Kyosuke Yamamoto[1], Hisashi Satoh[2], Masaru K. Nobu[3,8], Takashi Narihiro[1*]

[1]Bioproduction Research Institute, National Institute of Advanced Industrial Science and Technology (AIST), 2-17-2-1 Tsukisamu-Higashi, Toyohira-ku, Sapporo, Hokkaido, 062-8517 Japan
[2]Division of Environmental Engineering, Faculty of Engineering, Hokkaido University, North-13, West-8, Hokkaido, 060-8628 Japan
[3]Bioproduction Research Institute, National Institute of Advanced Industrial Science and Technology (AIST), Central 6, Higashi 1-1-1, Tsukuba, Ibaraki 305-8566, Japan
[4]Department of Civil and Environmental Engineering, National Institute of Technology, Anan College, 265 Aoki Minobayashi, Anan, Tokushima 774-0017, Japan
[5]Department of Civil and Environmental Engineering, Graduate School of Engineering, Tohoku University, 6-6-06 Aza-Aoba, Aramaki, Aoba-ku, Sendai, Miyagi 980-8579, Japan
[6]Department of Frontier Sciences for Advanced Environment, Graduate School of Environmental Studies, Tohoku University, 6-6-06 Aza-Aoba, Aramaki, Aoba-ku, Sendai, Miyagi 980-8579, Japan
[7]Department of Chemical Science and Engineering, National Institute of Technology, Miyakonojo College, 473-1 Yoshio-cho, Miyakonojo, Miyazaki, 885-8567, Japan
[8]Institute for Extra-cutting-edge Science and Technology Avant-garde Research (X-star), Japan Agency for Marine-Earth Science and Technology (JAMSTEC), 2-15 Natsushima-cho, Yokosuka, Kanagawa 237-0061, Japan

[¶]These authors contributed equally to this work.
[*]Co-corresponding authors:

### Materials and Methods

#### Enrichment culture experiments

For the experimental design of the 16S rRNA gene sequencing, metatranscriptome, and microscopy observations, we prepared seven parallel enrichment cultures (C-1, C-2, C-3, C-4, C-5, C-6, C-7), which were transferred 2 mL from the previous culture system C-d2-d1 on Day 33 (1, 2). One culture C-1 was used to check the cultivation condition of *Candidatus* Patescibacteria routinely (once per week) monitored by a microscope equipped with a phase-contrast apparatus (BX-53, Olympus, Japan). C2–C4 and C5–C7 were used to analyze the beginning (7 and 14 days) and end (21 and 31 days) of the culture phases, respectively. The inorganic basal media were prepared according to a previous study (3). The cultivation experiments were performed at 37 °C using 50 mL of serum vials containing 20 mL medium under $N_2/CO_2$ (80:20, v/v) atmosphere. These culture media contained 1 mM acetate, 0.1 mM adenosine 5'-monophosphate, uridine 5'-monophosphate, guanosine 5'-monophosphate, and cytidine 5'-monophosphate, 1% (w/v) MEM nonessential amino acids solution (cat no. 139-15651, FUJIFILM Wako Pure Chemical Co. Ltd., Tokyo, Japan), 1% (w/v) MEM essential amino acids solution (cat no. 132-15641, FUJIFILM Wako Pure Chemical Co. Ltd., Tokyo, Japan), *Methanothrix soehngenii* GP6 (DSM 3671, 0.2 mL/20 mL-medium), and *Methanosarcina barkeri* MS (DSM 800, 0.2 mL/20 mL-medium). *Methanothrix soehngenii* GP6 (DSM 3671) was precultivated at 37 °C for 4 weeks using 60 mM acetate and 10 mM potassium bicarbonate as substrates. *Methanosarcina barkeri* MS (DSM 800) was precultivated at 37 °C for 1 week using 10 mM methanol, 0.03% yeast extract (w/v), and 10 mM acetate as substrates. The biogas compositions ($CH_4$, $CO_2$, $N_2$, and $H_2$) of all cultures were determined by gas chromatography (Shimadzu, GC-8A, Japan) with a thermal conductivity detector fitted with a SHINCARBON-ST 50/80 stainless steel Column 4.0 m × 3.0 mm (ID) according to a previous study (4). On Days 14 and 31, approximately 17 mL of cultivated microorganisms was collected from the culture systems C-2–C-4 and C5–C-7 for RNA extraction. For DNA extraction and fluorescence *in situ* hybridization (FISH), approximately 1–3 mL of cultivated microorganisms in culture systems C1, C2–C4, and C5–C7 were sampled on Days 7, 14, 21, and 31, 7 and 14, and 21 and 31, respectively. Collected microbial cells were centrifuged at 17,750 g and stored at -80 °C until DNA/RNA extraction.

#### 16S rRNA gene sequence analysis

DNA was extracted from microbial cells using a FastDNA Spin Kit for Soil (MP Biomedicals, Santa Ana, California, USA) according to the manufacturer's protocol. The 16S rRNA genes were amplified using Univ515F–Univ909R according to a previous study (5, 6). The PCR products were purified using a QIAquick PCR purification kit (Qiagen, Valencia, CA, USA) according to the manufacturer's protocol. Sequence analysis was performed using the MiSeq Reagent kit v3 and MiSeq system (Illumina, San Diego, CA, USA). Raw 16S rRNA gene sequences were analyzed using QIIME 2 ver. 2021.4 (7) according to a previous study (5, 6). The 16S rRNA gene sequences were clustered by ≥ 97% similarity to operational taxonomic units (OTUs) using vsearch software (8). Taxonomic classification was carried out using classify-sklearn with the SILVA database version 138 (9).

#### Phylogenetic analysis

The phylogenetic tree of nearly full-length 16S rRNA gene sequences of *Ca.* Yanofskybacteria (LC715099) and 32-520 (LC715109 and LC715100) was constructed based on neighbor-joining methods in ARB version 7.0 (10) using the SILVA138.1 database for small subunit rRNA gene sequences. The sequences between *E. coli* positions 220 and 1460 were used for phylogenetic tree construction. A genome tree of previously reconstructed metagenome-assembled bins (2, 11) of *Ca.* Yanofskybacteria/UBA5738 PMX_810_sub (DRZ078160) and

32-520/UBA5633 PMX.108 (DRZ078158) and PMX.50 (DRZ078159) was constructed using IQ-TREE version 2.1.4-beta (-B 1000) with an automatically optimized substitution model (Q.yeast+F+R9) (12).

**Prediction of protein structures**

The protein structures of each gene were predicted using AlphaFold2 (13) and visualized using Chimera software version 1.15 (14). Furthermore, to elucidate the functions associated with these genes, the amino acid sequences were queried against the InterPro database (http://www.ebi.ac.uk/interpro/) to identify the functional domains. Subsequently, each protein structure predicted by AlphaFold2 and each identified domain, namely, the peptidoglycan binding domain (PDB: 1lbu), Ig-like domain (PDB: 6dlh), polycystic kidney disease domain (PDB: 2y72), galactose-binding-like domain (PDB: 3w5m), and thioredoxin-like domain (PDB: 4i5q), were superimposed using Chimera software.

**Metatranscriptomic analysis**

RNA was extracted from microbial cells using acid-chloroform, precipitated by cold ethanol, and purified by DNase treatment according to previous studies (15, 16). The total RNA concentration was quantified using a Quantus Fluorometer and QuantiFluor RNA system (Promega). The quality of the extracted RNA was confirmed using a 5200 Fragment Analyzer System and an Agilent HS RNA Kit (Agilent Technologies). The expression sequence data of one culture (C-5) on Day 31 were excluded from the following analysis due to the low RNA quality number (RQN=5.1) after RNA extraction, while other RNAs extracted from five cultures were ≥7.4 RQN. The rRNA was removed using riboPool (siTOOLs Biotech), and libraries were prepared using the MGIEasy RNA Directional Library Prep Set (MGI Tech Co., Ltd.) according to the manufacturer's protocol. The quality of the prepared libraries was checked using a Fragment Analyzer and dsDNA 915 Reagent Kit (Advanced Analytical Technologies). DNA circularization was performed using an MGIEasy Circularization Kit (MGI Tech Co., Ltd.), and then DNA nanoballs (DNBs) were generated using a DNBSEQ-G400RS High-throughput Sequencing Kit (MGI Tech Co., Ltd.). DNA sequencing (200 bp paired-end) was conducted using DNBSEQ-G400 (MGI Tech Co., Ltd.). The generated raw reads were trimmed using Trimmomatic 0.39 (SLIDINGWINDOW:6:30 MINLEN:50 LEADING:3 TRAILING:3 ILLUMINACLIP:trim.fa:2:30:10) (17). The trimmed reads were mapped to previously reconstructed metagenome-assembled bins (2, 11) of *Ca.* Yanofskybacteria/UBA5738 PMX_810_sub (DRZ078160), 32-520/UBA5633 PMX.108 (DRZ078158) and PMX.50 (DRZ078159), *Methanothrix* PMX.81 (DRZ078097), PMX.12 (DRZ078096), and PMX.35 (DRZ078098), and *Methanospirillum* PMX.141_sub (DRZ078095) using BBMap of the BBTools package v38.88 using a 99% similarity cutoff (minid = 0.99) (https://sourceforge.net/projects/bbmap/). The BBMap-predicted mapped reads per kilobase of transcript per million mapped reads (RPKM) values were normalized to $\log_2$(RPKM+1) values.

**Fluorescence *in situ* hybridization**

The collected samples were fixed with 4% paraformaldehyde in phosphate-buffered saline (PBS) for 4 hours at 4 °C and stored in 50% ethanol with PBS at -20 °C. FISH was performed as described previously (18). The hybridization and washing slides were incubated at 46 °C for 4–17 hours and 48 °C for 20 mins, respectively. An equimolar mixture (defined as EUB338mix) of EUB338 (5'- GCTGCCTCCCGTAGGAGT-3') (19), EUB338I (5'- GCAGCCTCCCGTAGGAGT-3'), EUB338II (5'- GCAGCCACCCGTAGGTGT-3'), and EUB338III (5'- GCTGCCACCCGTAGGTGT-3') (20) was used for the detection of all bacteria. Formamide concentrations used in this study were as follows: EUB338mix, 10%; ARC915 (5'-GTGCTCCCCCGCCAATTCCT-3') for all archaea (21), 35%; MX825 (5'-

TCGCACCGTGGCCGACACCTAGC-3’) for genus *Methanothrix* (21), 20%; Pac_683 (5’-TCAACGGATTTCACCCCTACAC-3’) for *Ca.* Yanofskybacteria (22) and MG1200 (5’-CGGATAATTCGGGGCATGCTG -3’) for order Methanomicrobiales (21), 25%; and previously designed 32-520-1066 probe (5’- GAGCAACTCAAGCCACCTGCTG -3’) for 32-520 lineages (11), 30%. PAC_683 shows a perfect match for the predominant order *Ca.* Yanofskybacteria OTU0011 (UBA5738) and the full length of the 16S rRNA gene sequence (100% similarity with OTU0011) and no perfect match for other detected *Ca.* Patescibacteria in all culture systems, which were confirmed by the ARB software package version 7.0 with the SILVA 138.1 database (10) (Fig. S2A). The 32-520-1066 probe covers 95.7% (22/23 match) of 32-520 lineages as a perfect match based on the SILVA138.1 database using TestProbe 3.0 [https://www.arb-silva.de/search/testprobe/] and perfectly matches the 16S rRNA gene sequences of targeted 32-520 (LC715109 and LC715100) in the cultures (Fig. S2A). In addition, the designed probe shows no perfect match for other detected *Ca.* Patescibacteria in all culture systems, which was also confirmed by ARB software with the SILVA138.1 database (10). All probes were labeled with FITC or Cy3. The FISH samples were also stained with 4’,6-diamidino-2-phenylindole dihydrochloride (DAPI). When double staining of 32-520 and *Methanospirillum* was performed, the fluorescence of the FITC-labeled MG-1200 probe disappeared, whereas single staining with the 32-520-1066 probe was successfully conducted. Although the reason for this phenomenon remains unclear, we used 32-520-1066 and ARC915 probes for the double staining of 32-520 and *Methanospirillum*, respectively. The microscopy images were observed by epifluorescence microscopy (BX-53, Olympus, Japan) with a color CCD camera (DP-74, Olympus, Japan). Phase-contrast, FISH, and DAPI micrographic images were uniformly processed across the entire images using Adjust color of Preview application on Mac OS 11.7.6 and ImageJ 1.53k (23). The observed FISH images were converted to 8-bit grayscale images, and then the brightness of the methanogenic archaeal cells was measured using ImageJ. For the *Methanothrix* cells attached to *Ca.* Patescibacteria, clear, weak, and no fluorescence was measured. The FISH brightness of *Methanospirillum* was measured from both the edges and center of the cells. Significant differences in the counted data were calculated using Tukey’s HSD test and Welch’s t test using R software version 4.1.0.

**Transmission electron microscopy**

On cultivation Day 33 of culture system C-1, approximately 1 mL of culture media was sampled and sandwiched with the copper disks in liquid propane at -175 °C. After the samples were frozen, the liquid was freeze-substituted with 2% glutaraldehyde, 1% tannic acid in ethanol, and 2% distilled water at -80 °C for 2 days. Dehydration was performed with anhydrous ethanol 3 times for 30 mins each. Infiltration was carried out by propylene oxide twice for 30 mins each, and the sample was put into a ratio of 7:3 mixtures of propylene oxide and resin for 1 hour. After volatilization of propylene oxide, the sample was transferred to fresh 100% resin and polymerized at 60 °C for 48 hours. Ultrathin sections at 70 nm were cut with a diamond knife using an ultramicrotome (Ultracut UCT, Leica, Vienna, Austria). The sections were stained with 2% uranyl acetate at room temperature for 15 mins and washed with distilled water, followed by secondary staining with lead stain solutions (Sigma–Aldrich Co., Tokyo, Japan) at room temperature for 3 mins. The grids were observed by transmission electron microscopy (TEM) (JEM-1500Plus, JEOL Ltd., Tokyo, Japan) at 100 kV. Digital images were observed with a CCD camera (EM-14830RUBY2, JEOL Ltd., Tokyo, Japan).

**Scanning electron microscopy**

The dehydrated sample in the **transmission electron microscopy** section was used for the scanning electron microscopy (SEM) experiment. The sample was placed in a 5:5 mixture of ethanol and t-butyl alcohol for 1 hour at room temperature. The sample was transferred to a t-butyl alcohol solution 3 times for 30 mins each. Then, freeze-drying was performed at 4 °C. Electroconductive coating was conducted using an osmium plasma heater (NL-OPC80NS, Nippon Laser & Electronics Laboratory), and the coating thickness was 30 nm. The images were observed with a scanning electron microscope (JSM-7500F; JFOL Ltd., Tokyo, Japan) at 3 kV.

**Deposition of DNA sequence data**

The raw sequence data and binned metagenome data are available in the DDBJ Sequence Read Archive database (DRA013834, DRA014327, and DRA016851). The genomic data of *Ca.* Yanofskybacteria/UBA5738 PMX_810_sub (DRZ078160), 32-520/UBA5633 PMX.108 (DRZ078158) and PMX.50 (DRZ078159), *Methanothrix* PMX.81 (DRZ078097), PMX.12 (DRZ078096), and PMX.35 (DRZ078098), and *Methanospirillum* PMX.141_sub (DRZ078095) can be downloaded from the URL of the DDBJ DRA014327 database "https://ddbj.nig.ac.jp/public/ddbj_database/dra/fastq/DRA014/DRA014327/". The nearly full-length 16S rRNA gene sequences of *Ca.* Yanofskybacteria/UBA5738 (LC715099) and 32-520/UBA5633 PMX.108 (LC715100) and PMX.50 (LC715109) are available in the DDBJ/EMBL/GenBank databases.

### Results and Discussion

**Linear relationships between *Ca.* Patescibacteria and methanogens based on the relative abundance**

The 16S rRNA gene-based relative abundance of *Ca.* Patescibacteria lineages and the predicted host methanogens showed that significant linear relationships ($p < 0.05$) existed between *Ca.* Yanofskybacteria–*Methanothrix* OTU0004 (Fig. S1B) and 32-520 OTU0072–*Methanospirillum* OTU0025 (Fig. S1C), indicating that the tendency of the growth of *Ca.* Patescibacteria is similar to that of the hosts. On the other hand, 32-520 OTU0014 was stably present during cultivation with no significant relationships with *Methanospirillum* OTU0025 (Fig. S1A and Fig. S1D). Thus, the growth of the 32-520 lineage OTU0014 might be different from that of OTU0072.

**Metatranscriptome overview**

To confirm their interactions based on gene expression levels, we performed metatranscriptomics for the enrichment cultures on Days 14 (triplicate) and 31 (duplicate). A total of 6.0–10.8 Gb sequences were obtained and mapped to the previously reconstructed metagenome-assembled bins of *Ca.* Yanofskybacteria/UBA5738 (PMX_810_sub), 32-520/UBA5633 (PMX.108 and PMX.50, which belonged to MWCK01 and UBA5633 at the genus level in the GTDB database (24), respectively), *Methanothrix* (PMX.81, PMX.12, and PMX.35), and *Methanospirillum* (PMX.141_sub) (2, 11). Compared with the 16S rRNA gene-based relative abundance, the gene expression levels of *Ca.* Yanofskybacteria/UBA5738 and 32-520/UBA5633 were low (percentage of mapped reads to the genomes per total reads was <1%) (Table S1); therefore, relatively low mapped proportions of the expressed genes on the genomes were obtained (Table S2). In a previous transcriptome study of a coculture of Saccharibacteria *Nanosynbacter lyticus* strain TM7x and the host XH001, only 880,000 reads were mapped on the Saccharibacteria TM7x, while an average of 27,000,000 mapped reads per sample were obtained for the host XH001 (25), suggesting that the gene expression levels of *Ca.* Patescibacteria/CPR are much lower than their hosts. In the gene expression of the methanogens, there were no significant differences between cultivation Days 14 and 31, and unique patterns of expression in the different *Methanothrix* species were not found in this study (Table S3); thus, further refinement of the cocultures is required to elucidate the details of *Ca.* Patescibacteria symbiosis.

X, McLean JS. 2022. Transcriptome of Epibiont Saccharibacteria Nanosynbacter lyticus Strain TM7x During the Establishment of Symbiosis. J Bacteriol 204.
