## Supplementary Figures for "Microscopic and metatranscriptomic analyses revealed unique cross-domain symbiosis between *Candidatus* Patescibacteria/candidate phyla radiation (CPR) and methanogenic archaea in anaerobic ecosystems"

Kyohei Kuroda[1*¶], Meri Nakajima[1,2¶], Ryosuke Nakai[1], Yuga Hirakata[3], Shuka Kagemasa[4,5], Kengo Kubota[5,6], Taro Q.P. Noguchi[7], Kyosuke Yamamoto[1], Hisashi Satoh[2], Masaru K. Nobu[3,8], Takashi Narihiro[1*]

[1]Bioproduction Research Institute, National Institute of Advanced Industrial Science and Technology (AIST), 2-17-2-1 Tsukisamu-Higashi, Toyohira-ku, Sapporo, Hokkaido, 062-8517 Japan
[2]Division of Environmental Engineering, Faculty of Engineering, Hokkaido University, North-13, West-8, Hokkaido, 060-8628 Japan
[3]Bioproduction Research Institute, National Institute of Advanced Industrial Science and Technology (AIST), Central 6, Higashi 1-1-1, Tsukuba, Ibaraki 305-8566, Japan
[4]Department of Civil and Environmental Engineering, National Institute of Technology, Anan College, 265 Aoki Minobayashi, Anan, Tokushima 774-0017, Japan
[5]Department of Civil and Environmental Engineering, Graduate School of Engineering, Tohoku University, 6-6-06 Aza-Aoba, Aramaki, Aoba-ku, Sendai, Miyagi 980-8579, Japan
[6]Department of Frontier Sciences for Advanced Environment, Graduate School of Environmental Studies, Tohoku University, 6-6-06 Aza-Aoba, Aramaki, Aoba-ku, Sendai, Miyagi 980-8579, Japan
[7]Department of Chemical Science and Engineering, National Institute of Technology, Miyakonojo College, 473-1 Yoshio-cho, Miyakonojo, Miyazaki, 885-8567, Japan
[8]Institute for Extra-cutting-edge Science and Technology Avant-garde Research (X-star), Japan Agency for Marine-Earth Science and Technology (JAMSTEC), 2-15 Natsushima-cho, Yokosuka, Kanagawa 237-0061, Japan

[¶]These authors contributed equally to this work.
[*]Co-corresponding authors:

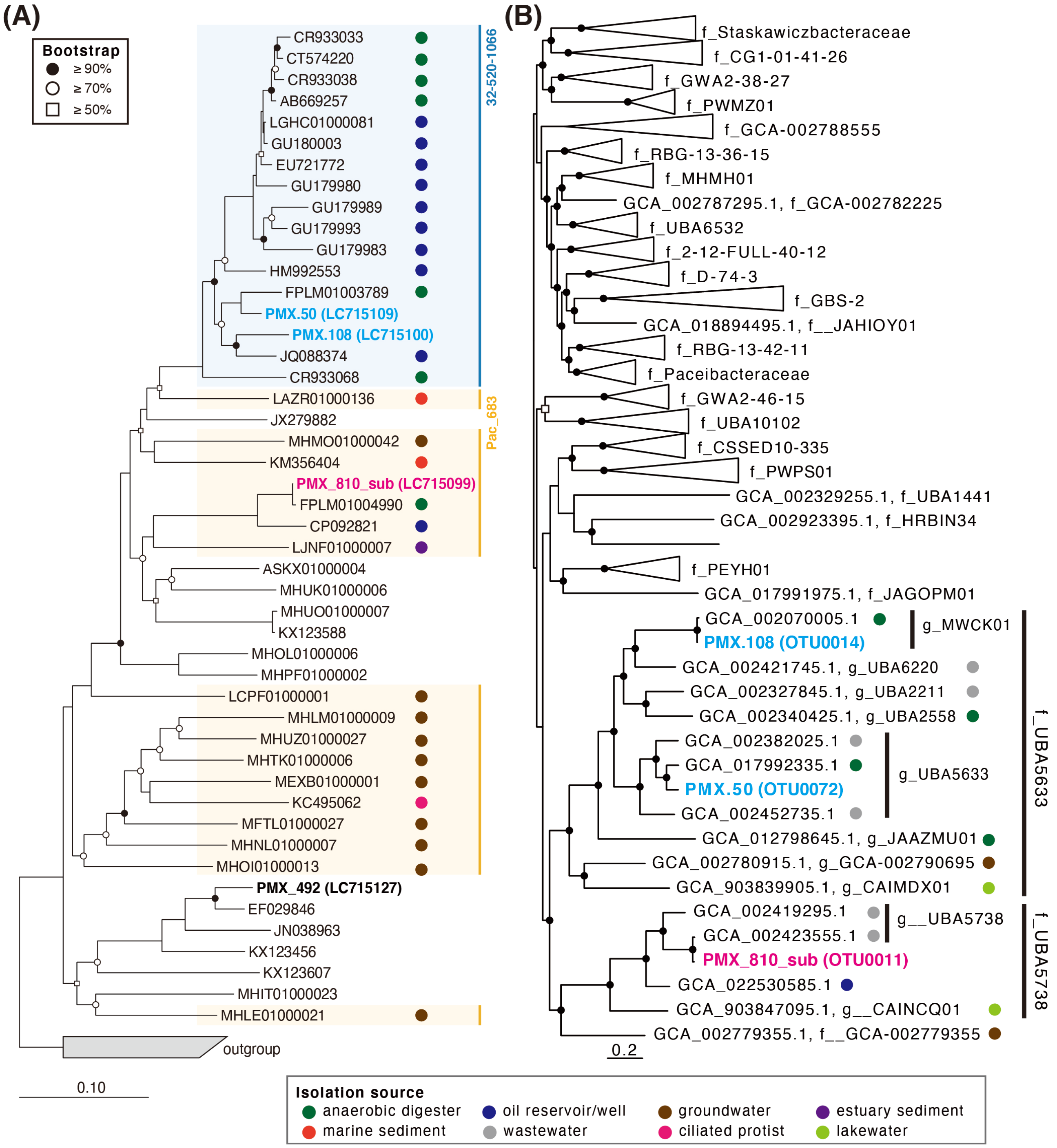


**Figure S1** Phylogenetic trees of order *Candidatus* Paceibacterales based on (A) 16S rRNA gene sequences and (B) concatenated phylogenetic marker genes of GTDBtk 2.0.0 (ver. r207) (15). The phylogenetic positions of the metagenomic bins PMX_810 and PMX.108/PMX.50 are shown in pink and blue, respectively. The 16S rRNA gene-based tree was constructed using the neighbor-joining method. Sequences that match the Pac_683 and 32-520-1066 probes are shown in yellow and blue layers, respectively.

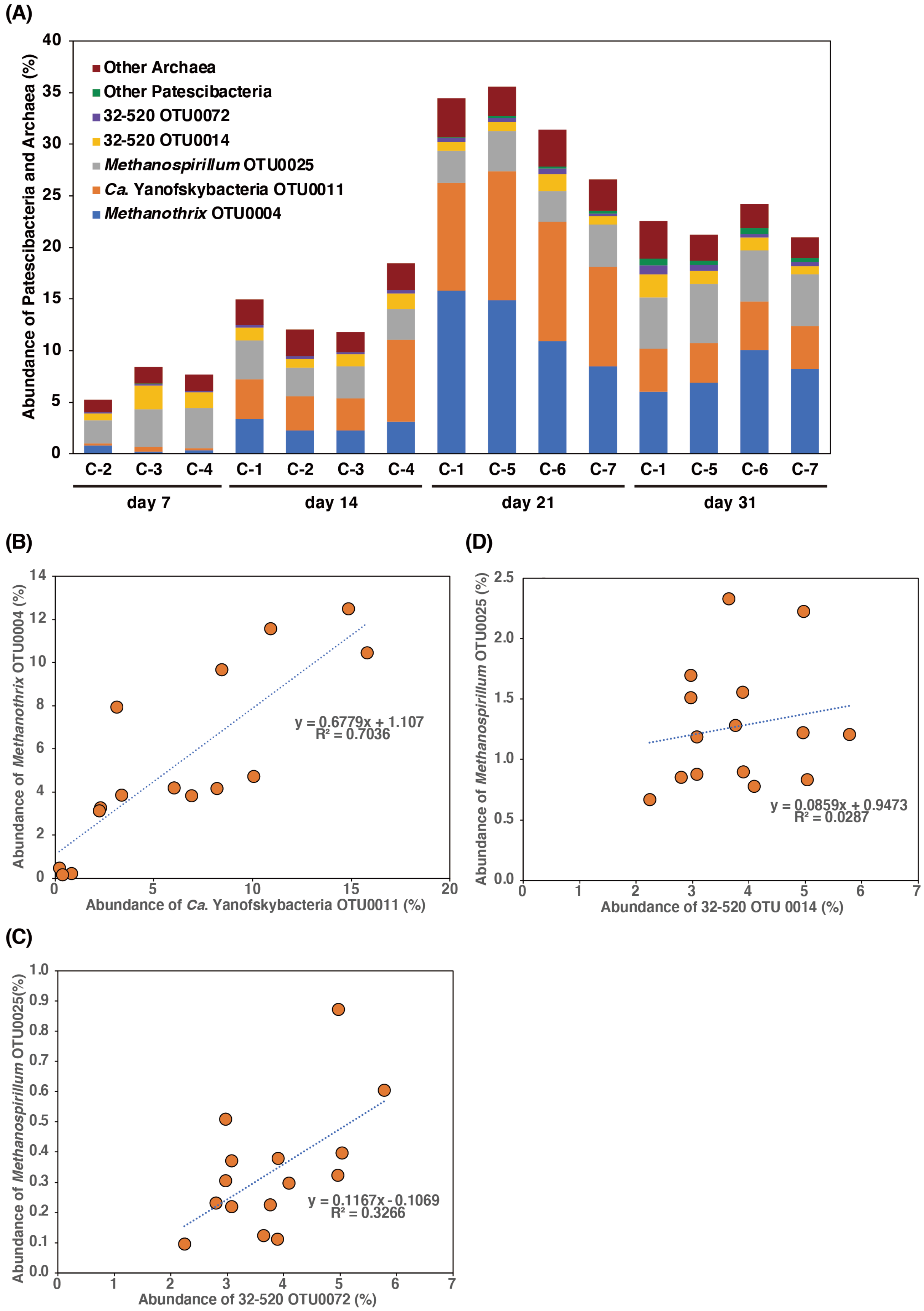


**Figure S2** (A) Relative abundance of predominant *Candidatus* Patescibacteria and methanogenic archaea in the culture systems based on 16S rRNA gene sequencing. (B)–(D) Linear regression analysis between predominant methanogens and *Ca*. Patescibacteria based on 16S rRNA gene-based relative abundance. (B) *Methanothrix* OTU0004 and *Ca*. Yanofskybacteria OTU0011, (C) *Methanospirillum* OTU0025 and 32-520 OTU0014, and (D) *Methanospirillum* OTU0025 and 32-520 OTU0072.

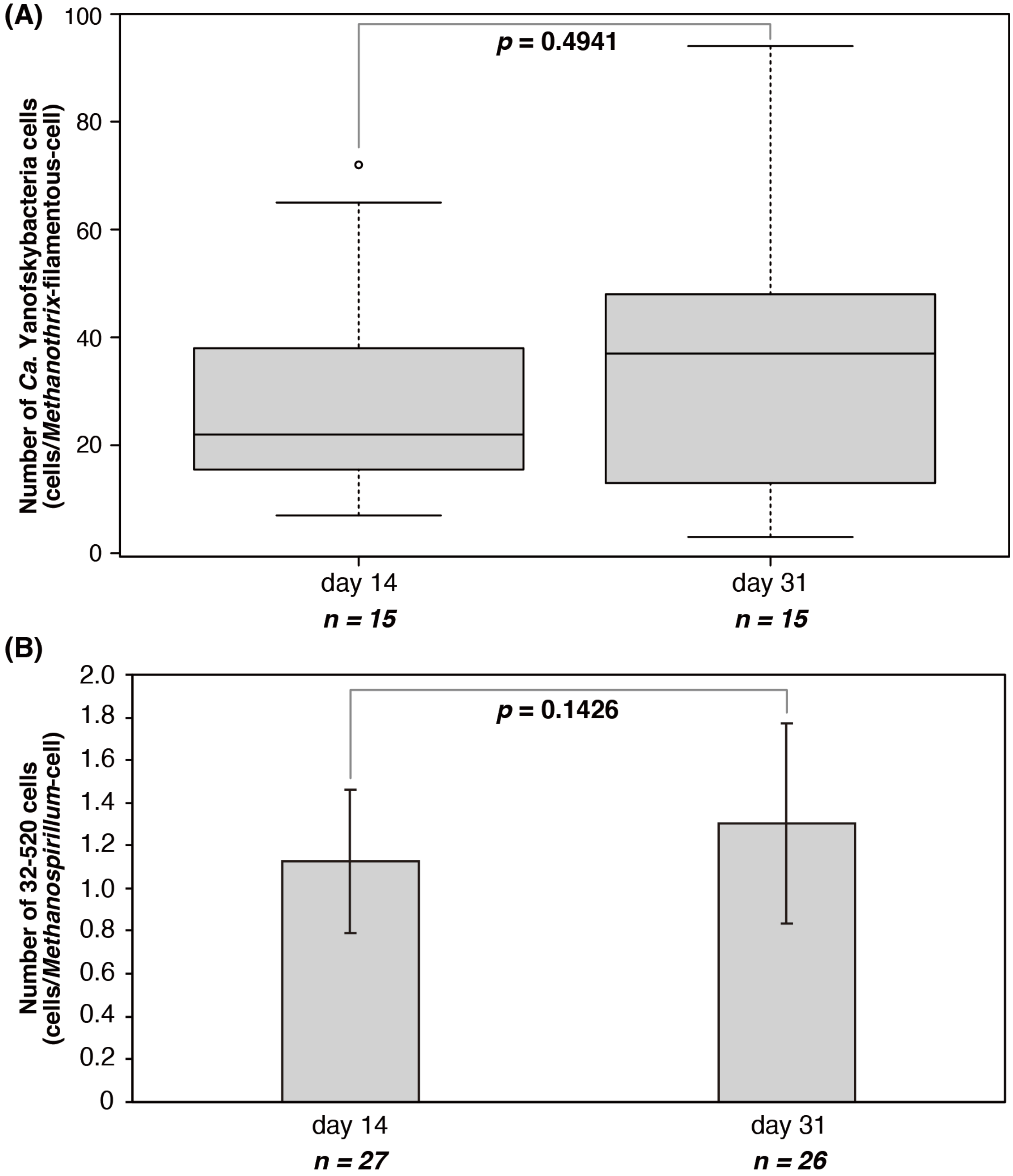


**Figure S3** (A) Number of *Candidatus* Yanofskybacteria cells attached to one *Methanothrix* filamentous cell on Days 14 and 31. (B) Number of 32-520 cells attached to one *Methanospirillum* cell on Days 14 and 31. The statistical analysis was performed based on Welch's t test.

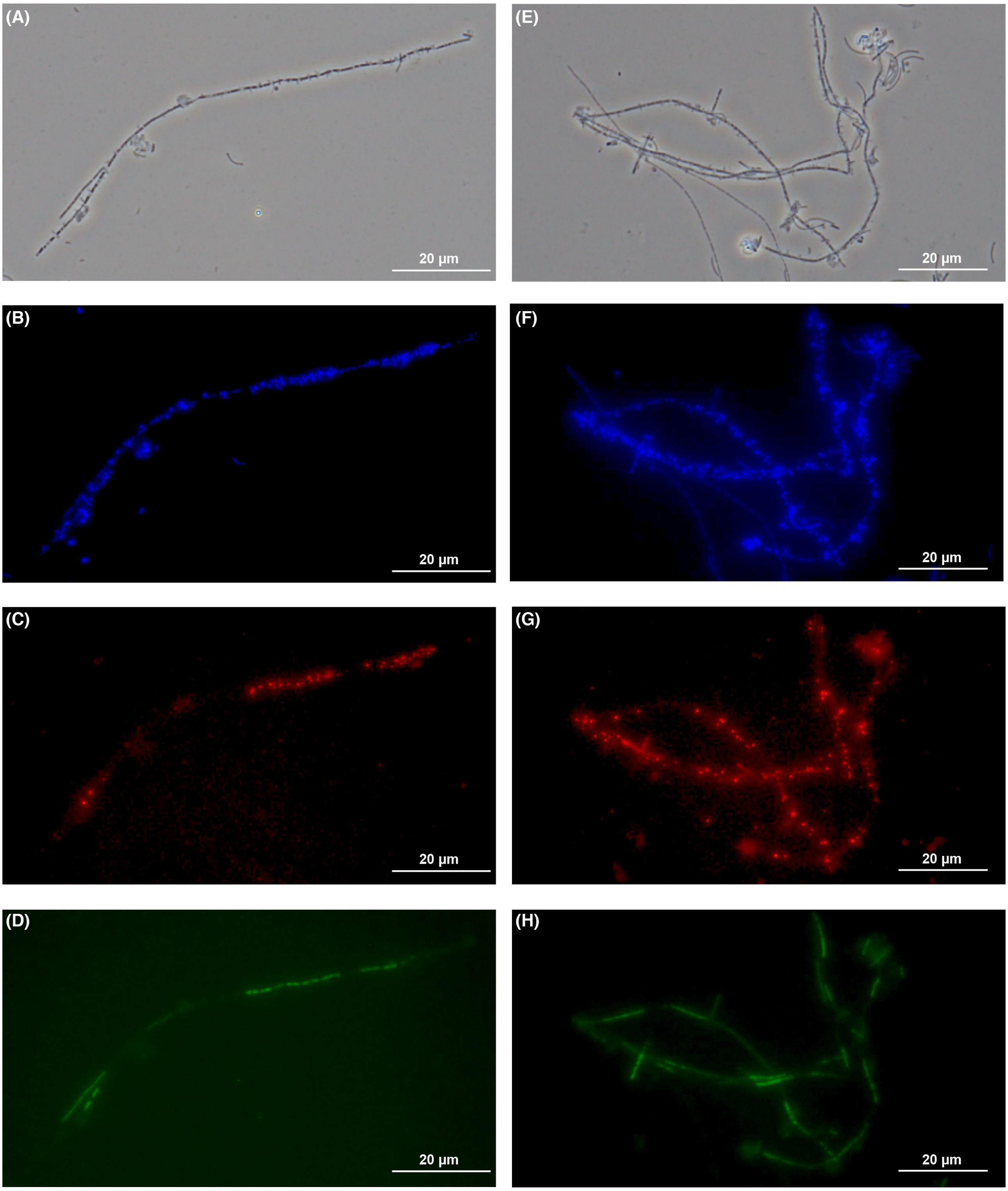


**Figure S4** Micrographs of (A) and (E) phase-contrast, (B) and (F) 4',6-diamidino-2-phenylindole dihydrochloride staining, (C), (D), (G), and (H) fluorescence *in situ* hybridization obtained from the culture system C-1 on (A)–(D) Days 14 and (G)–(H) 31. (C) and (G) *Ca.* Yanofskybacteria-targeting Pac_683-Cy3 probe and (D) and (H) *Methanothrix*-targeting MX825-FITC probe.

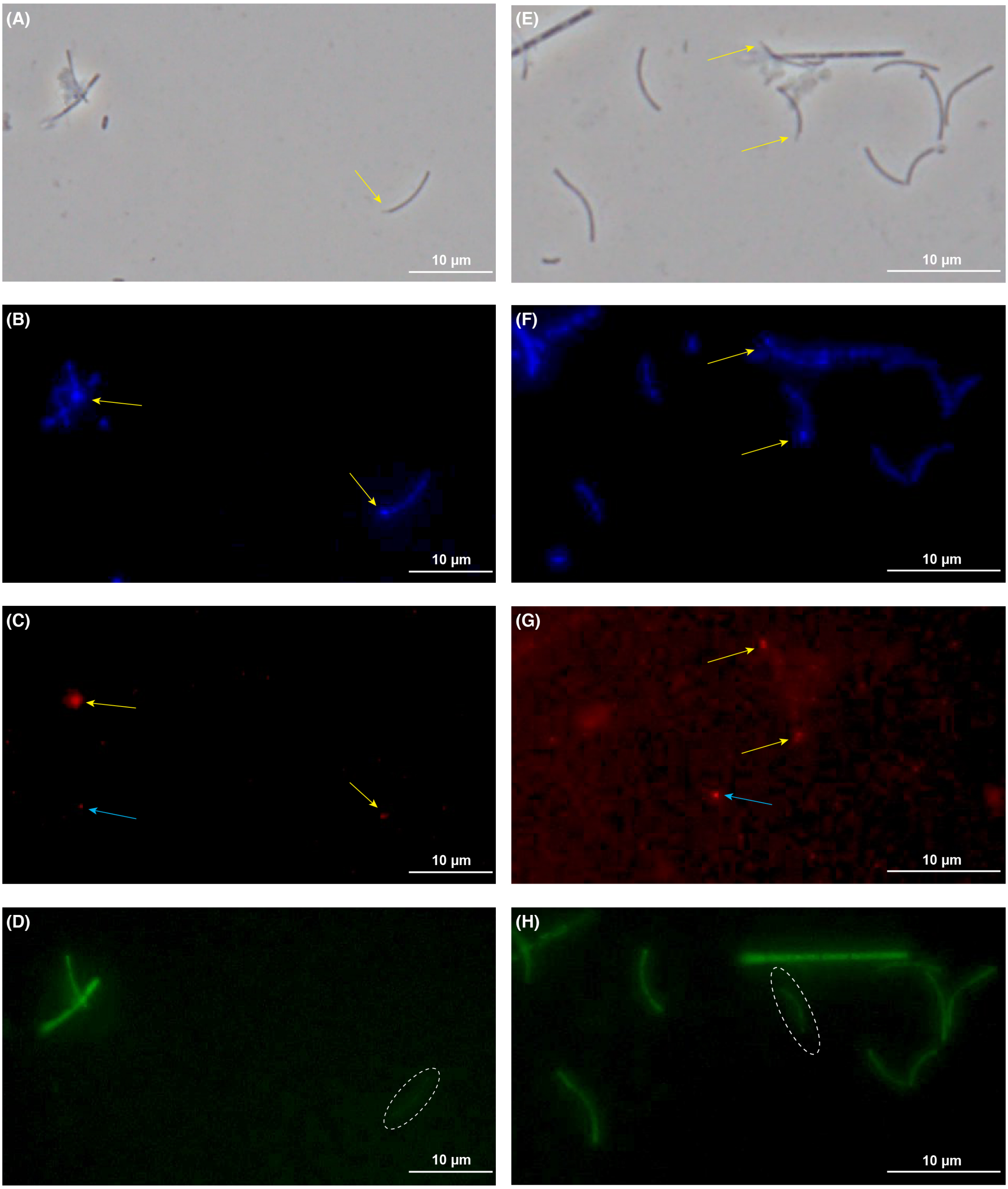


**Figure S5** Micrographs of (A) and (E) phase-contrast, (B) and (F) 4',6-diamidino-2-phenylindole dihydrochloride (DAPI) staining, (C), (D), (G), and (H) fluorescence *in situ* hybridization obtained from the culture system C-1 on (A)–(D) Days 14 and (G)–(H) 31. (C) and (G) 32-520-targeting 32-520-1066-Cy3 probe and (D) and (H) Archaea-targeting ARC915-FITC probe. Yellow arrows indicate FISH-detectable 32-520 cells. Light blue arrows indicate unspecific FISH signals that were not observed by phase-contrast and DAPI staining. Dashed white lines indicate weak or no FISH signals of *Methanospirillum*-like cells.
